# Optogenetic manipulation of medullary neurons in the locust optic lobe

**DOI:** 10.1101/334003

**Authors:** Hongxia Wang, Richard B. Dewell, Markus U. Ehrengruber, Eran Segev, Jacob Reimer, Michael L. Roukes, Fabrizio Gabbiani

## Abstract

Locust is a widely used animal model for studying sensory processing and its relation to behavior. Due to the lack of genomic information, genetic tools to manipulate neural circuits in locusts are not yet available. We examined whether Semliki Forest virus is suitable to mediate exogenous gene expression in neurons of the locust optic lobe. We subcloned a channelrhodopsin variant and the yellow fluorescent protein Venus into a Semliki Forest virus vector and injected the virus into the optic lobe of locusts (*Schistocerca americana*). Fluorescence was observed in all injected optic lobes. Most neurons that expressed the recombinant proteins were located in the first two neuropils of the optic lobe, the lamina and medulla. Extracellular recordings demonstrated that laser illumination increased the firing rate of medullary neurons expressing channelrhodopsin. The optogenetic activation of the medullary neurons also triggered firing of a postsynaptic, looming-sensitive neuron, the Lobula Giant Movement Detector (LGMD). These results indicate that Semliki Forest virus is efficient at mediating transient exogenous gene expression and provides a tool to manipulate neural circuits in the locust nervous system and likely other insects.

**New and Noteworthy:** Using Semliki Forest virus, we efficiently delivered channelrhodopsin into neurons of the locust optic lobe. We demonstrate that laser illumination increases the firing of the medullary neurons expressing channelrhodopsin and of an identified postsynaptic target neuron, the LGMD neuron. This technique allows to manipulate the neuronal activity in locust neural circuits using optogenetics.

## Introduction

Insects are widely used to address fundamental questions about brain mechanisms. Research on insects has broadened our knowledge and helped us understand the neural basis of complex behavior, e.g., communication and navigation in bees and ants (Evangelista et al., 2014; Srinivasan, 2010; Wehner, 2003), vision and motion detection in flies (Egelhaaf, 2008), olfactory learning and odor discrimination in flies, sphinx moths and locusts (Gupta and Stopfer, 2011), auditory processing in crickets (Göpfert and Hennig, 2016), as well as the mechanisms of neural development and genetics, most recently mainly in *Drosophila* (Hales et al., 2015; Spindler and Hartenstein, 2010). In these endeavors, genetic tools are helpful for dissecting neural circuits and deciphering the neural mechanisms underlying different behaviors. The Gal4-UAS system, for instance, is one of the most powerful ways of achieving targeted gene expression in *Drosophila* that has been adapted to other model systems (Asakawa and Kawakami, 2008; Brand and Perrimon, 1993; Busson and Pret, 2007; Imamura et al., 2003). This binary system is widely used to create transgenic flies by combining a driver (Gal4) and responder (UAS) line based on the properties of the yeast transcription factor GAL4 which activates its target genes by binding to UAS *cis*-regulatory sites. Since most neurons in *Drosophila* are small, they are unsuitable for intracellular dendritic recordings, making this model system of limited use for investigations of dendritic computations in single cells. On the other hand, insects with larger neurons such as locusts, crickets or moths have proven optimal for intracellular electrophysiological recordings. In most of these insects, however, it is not easy to manipulate gene expression and carry out genome editing due to lack of genome sequencing information and long generation times. Nevertheless, researchers have developed a variety of genetic tools for several such species. For example, *piggyBac*-derived cassettes have been integrated in the honeybee (*Apis mellifera*) expressing the fluorescent markers Rubia and EGFP under either an artificial or an endogenous promoter (Schulte et al., 2014). In another case, an odorant receptor co-receptor (orco) mutated ant germ line has been generated in *Ooceraea biroi* using CRISPR/Cas gene editing technology (Trible et al., 2017).

Locust is a popular model for studying behavior relying on visual motion, especially visually-evoked escape and collision-avoidance behavior (Fotowat and Gabbiani, 2011). Creating transgenic locust germ lines or developing an efficient neuronal transfection method in locusts would be desirable to increase the power of this model system. However, up to now, there are no known reports on any foreign transformation in the nervous system of locusts. Optogenetics is an efficient stimulation method to control neuronal activity using light-gated ion channels, such as channelrhodopsin, halorhodopsin and their variants (Arrenberg et al., 2009; Boyden et al., 2005; Ishizuka et al., 2006). It has been broadly used to map neural circuitry, study neuronal activity, control cardiac function, and treat photoreceptor degeneration and Parkinson disease (Adamantidis et al., 2007; Arenkiel et al., 2007; Bi et al., 2006; Gradinaru et al., 2009).

Semliki Forest virus (SFV) is an enveloped single-stranded, positive RNA virus, one of the members in the alphavirus family (Strauss and Strauss, 1994). In earlier work, wild-type SFV and mutant SFV A7(74) were used to drive LacZ and GFP expressions in pyramidal neurons of cultured hippocampal slices (Ehrengruber et al., 1999, 2003). Similarly, the less cytopathic mutant SFV(PD) (Lundstrom et al., 2003) drove protein expression in the rat calyx of Held *in vivo* (Wimmer et al., 2004). Since SFV is a mosquito-borne pathogen, it could possibly infect other non-host insect cells (Lwande et al., 2013), as has been shown for the Sindbis virus (Lewis et al., 1999). To test the possibility that SFV could drive foreign gene expression in locust neurons, we inserted into a SFV A7(74) based vector (Ehrengruber et al., 2003) the channelrhosopin variant, Chop-Wide Receiver (ChopWR), tagged with a fluorescent marker, Venus (Wang et al., 2009) and downstream of a strong ubiquitous promoter. ChopWR is a chimeric protein of Chop1 and Chop2 (Nagel et al., 2003), mediating a larger photocurrent (Wang et al., 2009). This plasmid was electroporated into baby hamster kidney 21 (BHK) cells for generating the virus, which was injected through the eye into the optic lobe of locusts.

In this paper, we show that viral replicons based on the SFV A7(74) strain successfully express ChopWR-Venus in medullary neurons of the locust optic lobe, enabling us to manipulate optogenetically the activity of medullary neurons and of the LGMD, a downstream neuron which plays a vital role in collision avoidance behavior (Fotowat and Gabbiani, 2011).

## Materials and Methods

### Generation of Semliki Forest virus with ChopWR-Venus and injection in locusts

The generation of SFV vectors was described in previous papers (e.g., Ehrengruber et al., 2011). Briefly, the ChopWR-Venus gene was first subcloned into the pENTR2B entry vector and then transferred to the pScaA7-RFA destination vector by using the attB1 and attB2 attachment sites through Gateway Technology (Thermo Fisher Scientific, Waltham, MA). The destination vector pScaA7-RFA was obtained from a modified SFV A7(74) vector plasmid, pSFV(A774nsP) (Ehrengruber et al., 2003), by moving A7(74)nsP1-4 into the pSCA plasmid (DiCiommo and Bremner, 1998). This plasmid uses the CMV/T7 promoters instead of the SP6 promoter and is compatible with Gateway Technology. The plasmid map of the pScaA7-RFA vector containing ChopWR-Venus is illustrated in Fig. 1A. The ChopWR-Venus gene was inserted downstream of the endogenous SFV subgenomic promoter that follows the sequence of SFV non-structural protein 4 (nsP4). Unique restriction sites are indicated in Fig. 1A, as is the simian virus 40 polyadenylation (SV40 polyA) terminator sequence, the Ampicillin resistance gene, and the pBR322 origin of replication. The resulting plasmid and the auxiliary plasmid pSFV-helper2 were purified and linearized with the restriction enzyme Spe I (New England Biolabs, Ipswich, MA; NEB). The pENTR2B and pScaA7 plasmids were a gift from Dr. Keith Murai (McGill University) and the pSFV-helper2 plasmid was a gift of Dr. Alan L. Goldin (University of California, Irvine). T7 and Sp6 RNA polymerase (Thermo Fisher Scientific) were used to catalyze the formation of RNAs from linearized pScaA7-ChopWR-Venus and pSFV-helper2 DNAs, respectively. To produce viruses, *in vitro* transcribed RNA from pScaA7-ChopWR-Venus and pSFV-Helper2 were co-electroporated into BHK-21 cells. After that, BHK-21 cells were incubated for 24-48 h in minimum essential (α-MEM) medium containing 5% fetal bovine serum (FBS) at 31 °C with 5% CO_2_ (Thermo Fisher Scientific, Waltham, MA). The SFV replicons were harvested and activated by 500 μg/ml α-chymotrypsin for 30 minutes at room temperature, and the reaction was stopped by 250 μg/ml aprotinin (Sigma-Aldrich, St.

**Figure 1.**
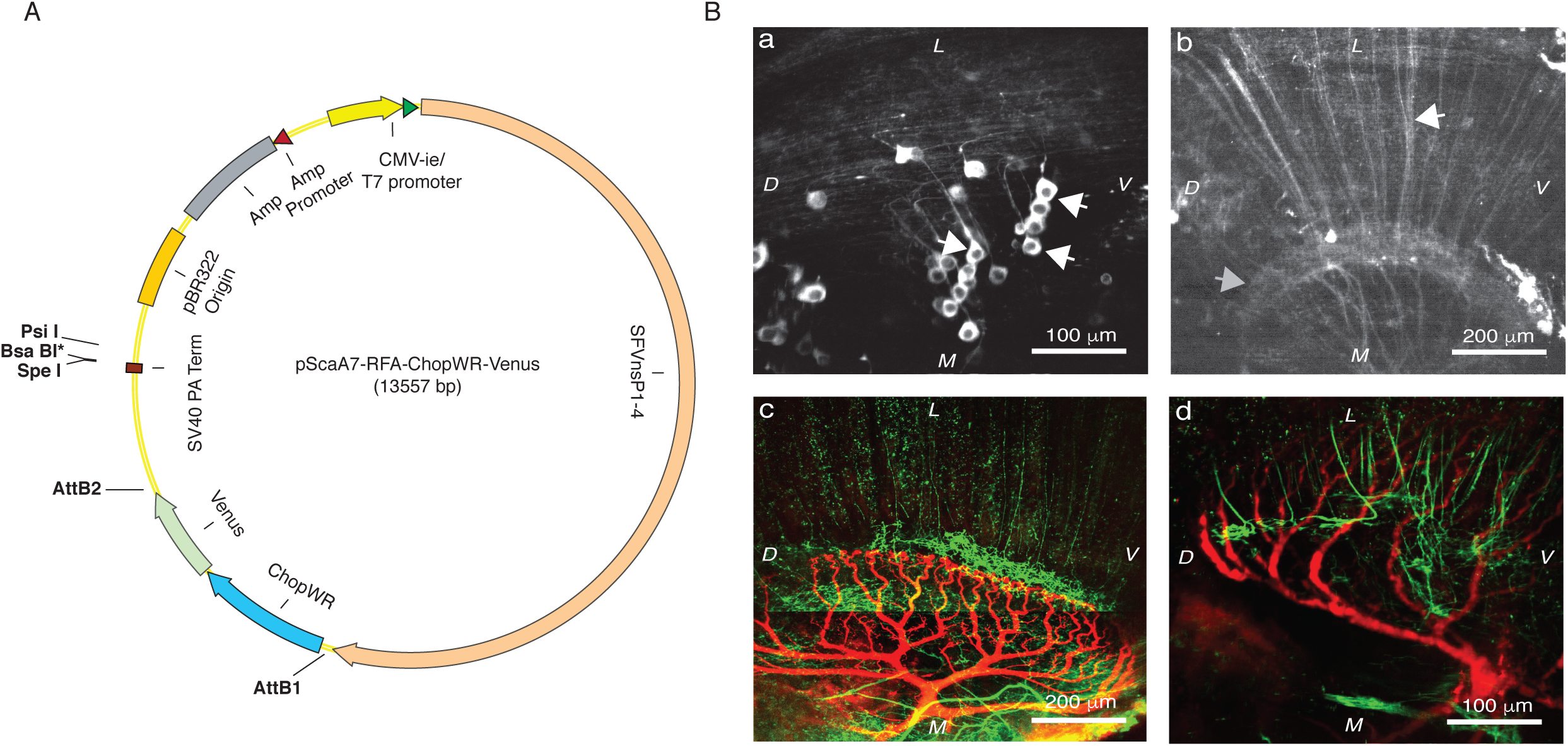
Semliki Forest Virus (SFV) drives Chop-wide receiver (ChopWR)-Venus expression in medullary neurons of the locust optic lobe. **A**, Schematic diagram showing the plasmid used to generate the SFV A7(74) based vectors encoding ChopWR-Venus. The viral backbone is derived from pSFV(A774nsP) (Ehrengruber, et al. 2003). **B**, a), Medullary neurons somata expressed ChopWR-Venus in the locust optic lobe (white arrows). b), Bundles of transmedullary axons expressing ChopWR-Venus (white arrow) travel towards the lobula neuropil (grey arrow). c) and d), Double stain of the LGMD excitatory dendritic field (Alexa 594, red) and Venus-labeled transmedullary neuron terminal arbors (green) show close apposition in the lobula. Abbreviations, *L*: lateral, *M*: medial, *D*: dorsal, *V*: ventral. Scale bars are listed in each panel.

Louis, MO). The SFV virus titers were ∼1×10^5^ infectious particles per ml. Experiments were done using both male and female locusts, *Schistocera americana*, 8-10 weeks old. The locusts were fed with grass sprayed with a solution containing *all-trans*-retinal (1 mM, Toronto Research Chemicals, North York, ON, Canada) for 2 days before being injected with ∼1-2 μl of viral solution into the right eye using a glass pipette under a Leica stereomicroscope. All protocols were approved by the Bio-Environmental Safety Committee of Baylor College of Medicine.

### Pilot experiments with other virus expression systems

In preliminary experiments, we tested several additional viral delivery vectors. Sindbis virus with the SP6 promoter was prepared similarly as described above. The titer for Sindbis virus was 5×10^6^ infectious particles per ml. Recombinant, GFP-tagged baculovirus with the polyhedrin promoter was purchased from a commercial supplier (#C14, AB Vector; titer: 10^8^ pfu/ml). Recombinant adeno-associated virus (AAV)-eGFP with the CMV promoter was a gift from Dr. Matthew Rasband (Baylor College of Medicine; titer: 1×10^13^ GC/ml). In each case, 1-2 μl of solution containing each virus was injected into the locust right eye using a glass pipette under a Leica stereomicroscope. Other procedures were as for the SFV vector.

### Electrophysiology

The dissection of the locust optic lobe was described in previous papers (e.g., Gabbiani et al., 2002). The LGMD was stained with Alexa 594 by intracellular negative current pulse injection. The Venus-tagged presynaptic neurons, especially their axon terminals, and the LGMD were visualized using two-photon microscopy. The excitation wavelength was set at 830 nm for Alexa 594 and 920 nm for Venus. Sharp electrodes (∼10-20 MΩ) were used for intracellular recording from the LGMD. Initially, spikes of the descending contralateral movement detector (DCMD) neuron were recorded extracellularly by positioning hook electrodes around the ventral nerve cord. DCMD spikes allowed us to identify the LGMD in the lobula since they are in one-to-one correspondence with LGMD spikes (O’Shea and Williams, 1974). A 488 nm Cyan Laser (Newport, Model No. PC13589, Ottawa, ON, Canada) with maximum output of 20 mW was used to stimulate ChopWR-expressing neurons in the optic lobe. Medullary neuronal activity was recorded by using a pair of 5 MΩ tungsten electrodes (FHC, Bowdoin, ME; see Wang et al., 2018 for details).

### Optogenetic stimulation

Optic fibers (Thorlabs, Newton, NJ) with diameters of 10, 25 and 200 μm were used to deliver laser light. The optic fiber was connected to the 488 nm Cyan laser via a collimator (F240FC-A, Thorlabs). The area of the incident laser beam arriving at the optic lobe was ∼1 mm^2^. The laser power was varied between 2 and 20 mW by inserting neutral density filters at the output port of the laser, immediately prior to the collimator, with transmission rates of 10%, 25%, 40%, 63% and 79%, respectively. To restrict the number of activated neurons, a custom-designed laser probe yielding a laser beam with a diameter of ∼10 μm was used in a subset of experiments (Segev et al., 2016). The time interval between two successive laser stimulations was 2 minutes to minimize desensitization of the responses. To minimize photoreceptor activation from reflected laser light, the eye was covered with black wax and/or black vinyl tape during laser stimulation. Despite this precaution, light hitting the back of the eye caused small, transient photoreceptor activations when the laser switched on and off.

### Injection of picrotoxin in the lobula

To investigate whether inhibitory neurons presynaptic to the LGMD had been activated by the laser stimulation, picrotoxin (5 mM; Sigma-Aldrich) dissolved in water was puffed along the dorsal edge of the lobula, close to the region where the inhibitory dendrites of the LGMD’s field C arborize. The injected solution contained 0.5 % of the colorant fast green (Sigma-Aldrich) to visualize the tip of the injection pipette and the amount of solution injected in the lobula. The injection pipette’s tip diameter varied between 1 and 2.5 μm. After injection, the dye diffused around the injection site and stayed confined to the lobula. A picospritzer was used to control the duration and puffing pressure (8 psi/55 kPa; WPI, Sarasota, FL). Based on earlier work (Dewell and Gabbiani, 2018), the estimated final concentration of drug at the level of field C was ≤ 200 μM.

### Data analysis and statistics

Custom Matlab (The MathWorks, Natick, MA) code was used for data analysis. The raw data recorded from medullary neurons were first normalized; the spikes were then detected with a set threshold (see Wang et al., 2018, for details). Spikes within 1.5 ms of a previously detected spike were excluded. To calculate instantaneous firing rates (IFRs) during looming stimuli, the spike train of the LGMD and the medullary neurons were convolved with a Gaussian filter that had a standard deviation of 20 ms. In Fig. 2E, the membrane potential (V_m_) of the LGMD was median filtered over a time window of 25 ms to eliminate spikes and reveal the subthreshold V_m_ time course. For the same reason, in Fig. 2F the average LGMD’s V_m_ during laser stimulation was calculated as its median value during the time interval when the laser was turned on. It was compared with the LGMD’s V_m_ during spontaneous activity, calculated as the median value during the time interval from the start of recording to the start of laser stimulation (>1 s). Medians for each trial where then averaged across 3-5 trials per animal. When using the custom laser probe no spiking was evoked and the LGMD’s V_m_ during laser stimulation was calculated as the mean of V_m_ during the time when the laser was turned on (Fig. 5B). It was compared with the LGMD’s V_m_ during spontaneous activity, calculated as the mean within 1 s before the start of laser stimulation (Fig. 5C).

**Figure 2.**
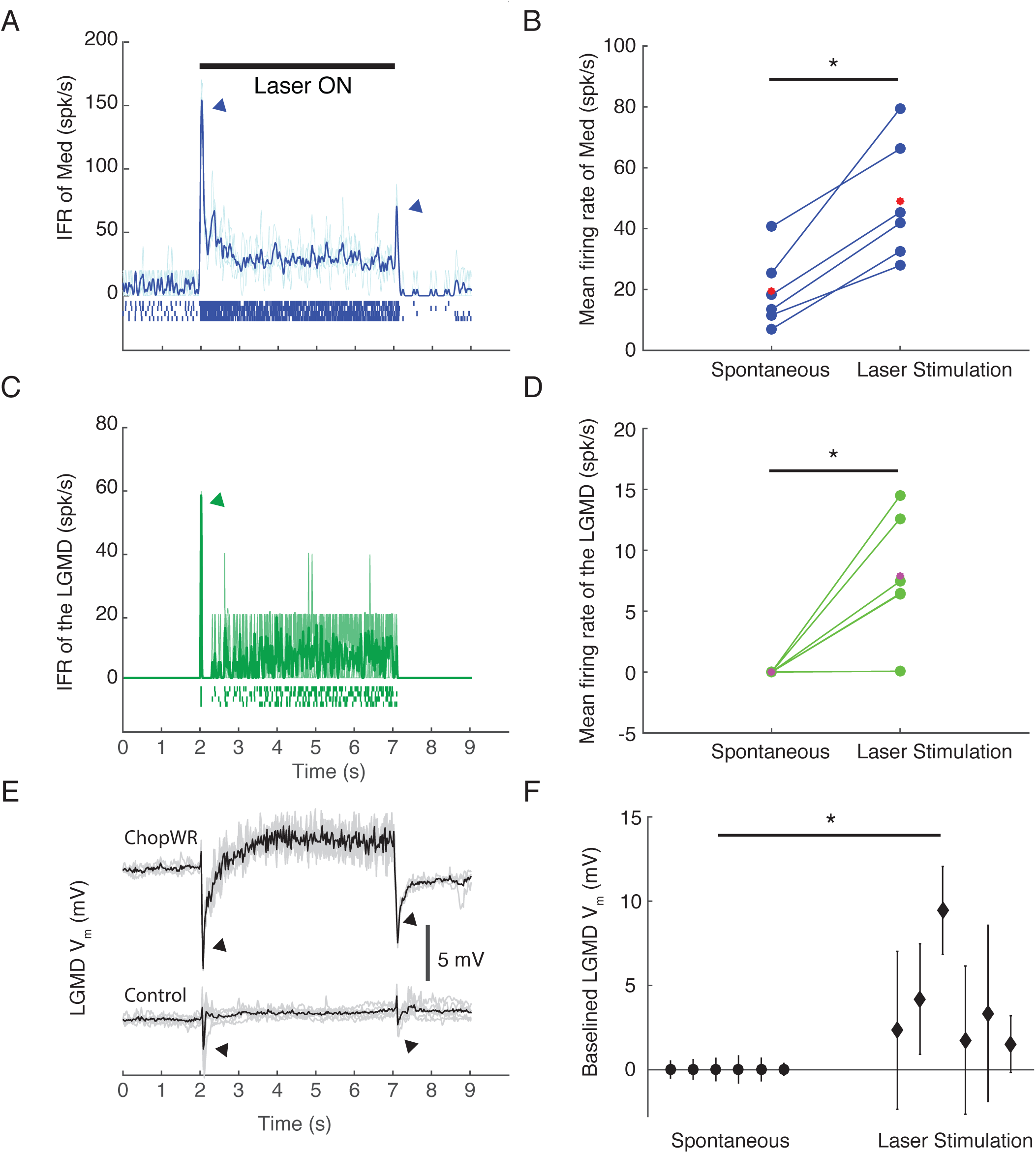
Laser stimulation via optic fiber with a diameter of 200 μm activated the medullary neurons (Med) expressing ChopWR-Venus and the LGMD. **A,** The instantaneous firing rate of the medullary neurons expressing ChopWR-Venus was increased during 5 s of 488 nm laser stimulation; top, laser stimulation timing; bottom, blue trace is the averaged firing rate across 4 trials (light blue traces). Rasters below the IFR show the medullary neuronal spikes. **B**, The mean firing rate of the medullary neurons expressing ChopWR-Venus across 6 locusts (red dots) was compared with and without laser stimulation; * indicates p = 0.0156 (one-sided WSRT). **C,** The instantaneous firing rate of the LGMD increased during 5 s of 488 nm laser stimulation; green trace is the averaged firing rate across 4 trials (light green traces). Rasters below the IFR show the LGMD spikes. **D**, The mean firing rate of the LGMD across 6 locusts (red dots) was compared with and without laser stimulation; * indicates p = 0.0156. **E**, The LGMD V_m_ was depolarized during 5 s of 488 nm laser stimulation in a ChopWR-expressing locust (top), while no depolarization was observed in an un-transfected control (bottom). Black trace represents the averaged V_m_ and gray traces are individual trials (4 trials in the ChopWR-expressing locust and 6 trials in the wild type locust). **F**, Plot of the mean median LGMD V_m_ (± 1 s.d.) with and without laser stimulation in 6 animals. * indicates p = 0.0156.

**Figure 5.**
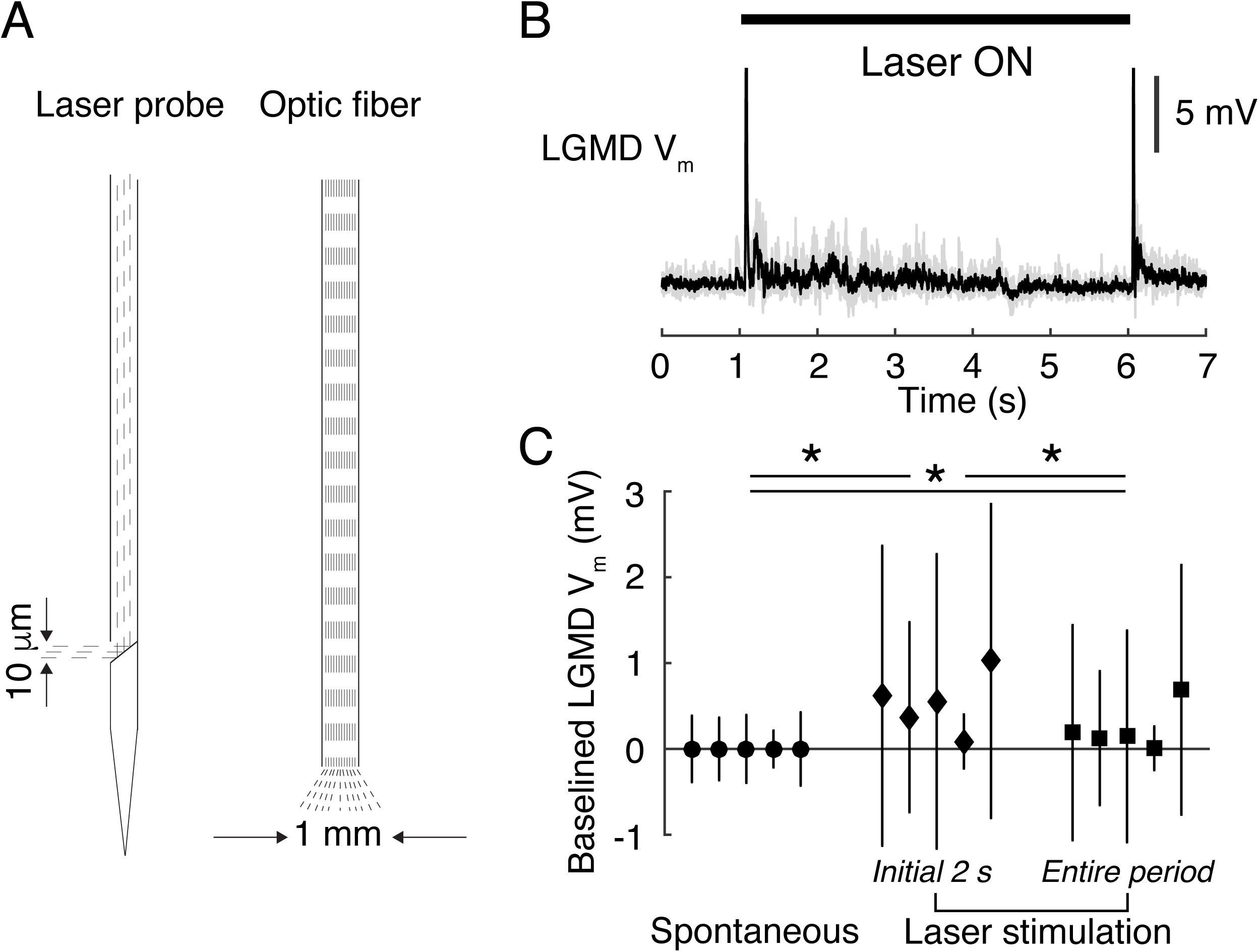
A laser probe narrowing the region activated by the laser elicited only EPSPs in the LGMD. **A**, Schematics of the laser probe (left) and optic fiber (right). Dashed lines indicate light path. In the laser probe, the beam exits perpendicular to the shaft thanks to a mirror. **B**, Laser probe stimulation (5 s, 488 nm) triggered EPSPs in the LGMD. Top, laser stimulation timing; bottom, the averaged LGMD V_m_ (black trace) across 4 trials (gray traces). **C**, Comparison of the mean across 3-5 trials of the LGMD V_m_ (± 1 s.d.) for 5 locusts with and without laser stimulation. For each animal the mean V_m_ was higher during the laser stimulation and higher during the first 2 s than the last 3 s of laser stimulation. * indicates p = 0.0312 by a one-sided WSRT.

The one-sided Wilcoxon signed-rank test (WSRT) was used to compare the statistical differences between groups of spontaneous activities and activities stimulated by optogenetics with or without picrotoxin treatment. All the data are described as mean ± s.d. (standard deviation).

## Results

### Semliki Forest Virus drives expression of ChopWR-Venus in locust medullary neurons

In pilot experiments, we tested with little success the capacity of several viruses to transfect locust optic lobes neurons, including adeno-associated virus (AAV), Sindbis and baculovirus. Although AAV is not known to infect arthropods, other members of its family, as well as Sindbis and baculovirus do (Cotmore et al., 2014; Lewis et al., 1999; Oppenheimer et al., 1999). In contrast, three days after recombinant SFV injection, ChopWR-Venus was observed expressing on the membrane of medullary neurons cell bodies, axonal fibers, and presynaptic terminals in the lobula (Fig. 1B, a-d). When the LGMD was concurrently stained with the fluorescent dye Alexa 594, some axon terminals overlapped with the dendritic branches of the LGMD (Fig. 1B, c, d). Additionally, stained neurons were also observed in the lamina in some experiments, when viral solution was deposited there upon retraction of the injection pipette (not shown). These results indicate that the SFV A7(74) plasmid vector can efficiently deliver a gene of interest into neurons of the medulla (and lamina) of the locust optic lobe. Five of 70 animals injected with virus died; all other locusts were healthy and did not appear to be affected negatively by the manipulation during the experiments.

### Optogenetic stimulation of medullary neurons activates the LGMD

As demonstrated in Fig. 2A, the instantaneous firing rate (IFR) of transfected medullary neurons increased in response to a 5 s long laser pulse. On average, the spontaneous firing rate of medullary units recorded from a pair of tungsten electrodes was 19.4 ± 12.2 spk/s (mean ± s.d.), while optogenetics stimulation increased the rate to 48.9 ± 20.0 spk/s (Fig. 2B). The IFR of the LGMD increased as well (Fig. 2C). On average, the mean firing rate of the LGMD in response to laser stimulation increased from 0 to 7.9 ± 5.1 spk/s (Fig. 2D). The turning ON and OFF of the laser caused brief spike bursts in the medullary neurons (Fig. 2A, arrowheads). Correspondingly, the LGMD fired an initial spike right after the ON transition (Fig. 2C, arrowhead) which was immediately followed by a transient membrane potential (V_m_) hyperpolarization of ∼1 s duration, also observed right after the laser was turned OFF (Fig. 2E top, arrowheads). During the laser stimulation, the LGMD V_m_ was depolarized by 3.8 ± 3.0 mV (Fig. 2E, F) in the ChopWR expressing locusts. In uninjected controls, transient responses occurred with laser onset and offset, but no sustained membrane potential depolarization was observed (Fig. 2E bottom). These results imply that optogenetic manipulation of medullary neurons is able to modulate the activity of one downstream target neuron, the LGMD.

### Block of inhibition enhances LGMD firing to optogenetic stimulation

To isolate the excitatory inputs to the LGMD, the GABA_A_ receptor antagonist, picrotoxin was puffed at the dorsal edge of the lobula, where inhibitory dendritic branches of the LGMD are located. Compared with the control group, block of inhibition increased the firing of the LGMD in response to the laser stimulation (Fig. 3A and B). The LGMD firing rate caused by the laser stimulation increased from 1.8 ± 0.8 to 7.9 ± 4.3 spk/s after adding picrotoxin (Fig. 3C). Addition of picrotoxin removed the hyperpolarizations observed at the onset and offset of the laser pulse (Fig. 2E), and instead the luminance change caused by the laser turning on and off produced transient bursts with peak firing rates of 69.9 ± 75.1 and 48.8 ± 48.2 spk/s after GABA_A_ blockade (Fig. 3B, D and E). These results demonstrate that the combination of blocker and optogenetic stimulation was effective at isolating the excitatory input to the LGMD.

**Figure 3.**
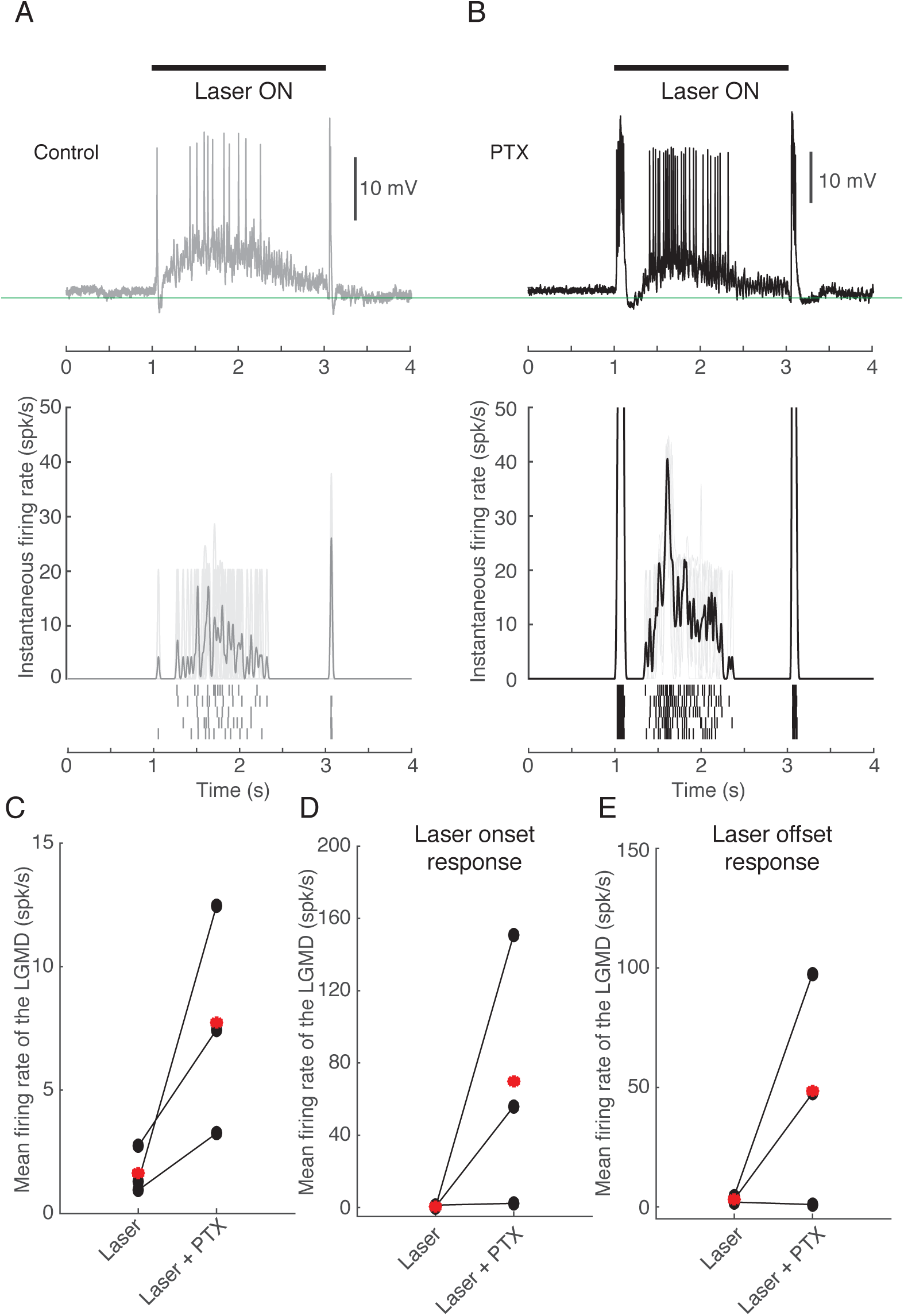
Laser stimulation triggers inhibitory inputs to the LGMD that can be blocked by the GABA_A_ receptor antagonist picrotoxin (PTX). **A**, Laser stimulation (2 s, 488 nm) via an optic fiber with diameter of 10 μm triggered the firing of the LGMD. Top, laser stimulation timing; middle, the LGMD V_m_ from 1 trial; bottom, the averaged LGMD IFR (gray trace) across 5 trials (light gray traces). The rasters below are the spikes of the LGMD from 5 trials. **B**, Puffing PTX increased laser-triggered firing in the LGMD. Top, laser stimulation timing; middle, the LGMD V_m_ from 1 trial; bottom, the averaged LGMD IFR (gray trace) across 5 trials (light gray traces). The rasters below are the spikes of the LGMD from 5 trials. **C**, The mean firing rate of the LGMD triggered by laser stimulation was compared with and without puffing PTX. **D** and **E**, mean firing rate of the LGMD triggered by laser onset (D) and offset (E) were compared with and without puffing PTX. Red symbols indicated the mean values across 3 locusts.

### Laser power affects the firing of the LGMD

Next, we tested whether optogenetic activation of the LGMD depends on the strength of the laser power stimulus used to activate channelrhodopsin. As demonstrated for one example in Fig. 4A, the mean number of spikes of the LGMD across 5 trials increased from 19.8 ± 3.3 to 44.6 ± 14.7 when power increased from 2 to 8 mW. However, at the higher power of 16 mW, the number of LGMD spikes elicited by the laser stimulus was slightly lower, 33.2 ± 11.9. We further investigated the effect of laser power in 5 animals by using 6 values varying from 2 to 20 mW (Fig. 4B). As shown in Fig. 4B, we found that spiking increased when power was increased from 2 to 5 and 8 mW (18.4 ± 6.2, 31.3 ± 19.1 and 37.7 ± 23.3, respectively). However, higher powers of 13, 16 and 20 mW resulted in slightly decreased spiking output than at 8 mW (28.2 ± 19.0, 28.4 ± 16.9 and 30.6 ± 17.4, respectively). The decrement might be caused by desensitization of channel rhodopsin at stronger laser power (Lin, 2011; Wang et al., 2009). These results indicate that optogenetic activation of the LGMD can be modulated by the laser power strength for a given expression level of Chop-WR in medullary neurons.

**Figure 4.**
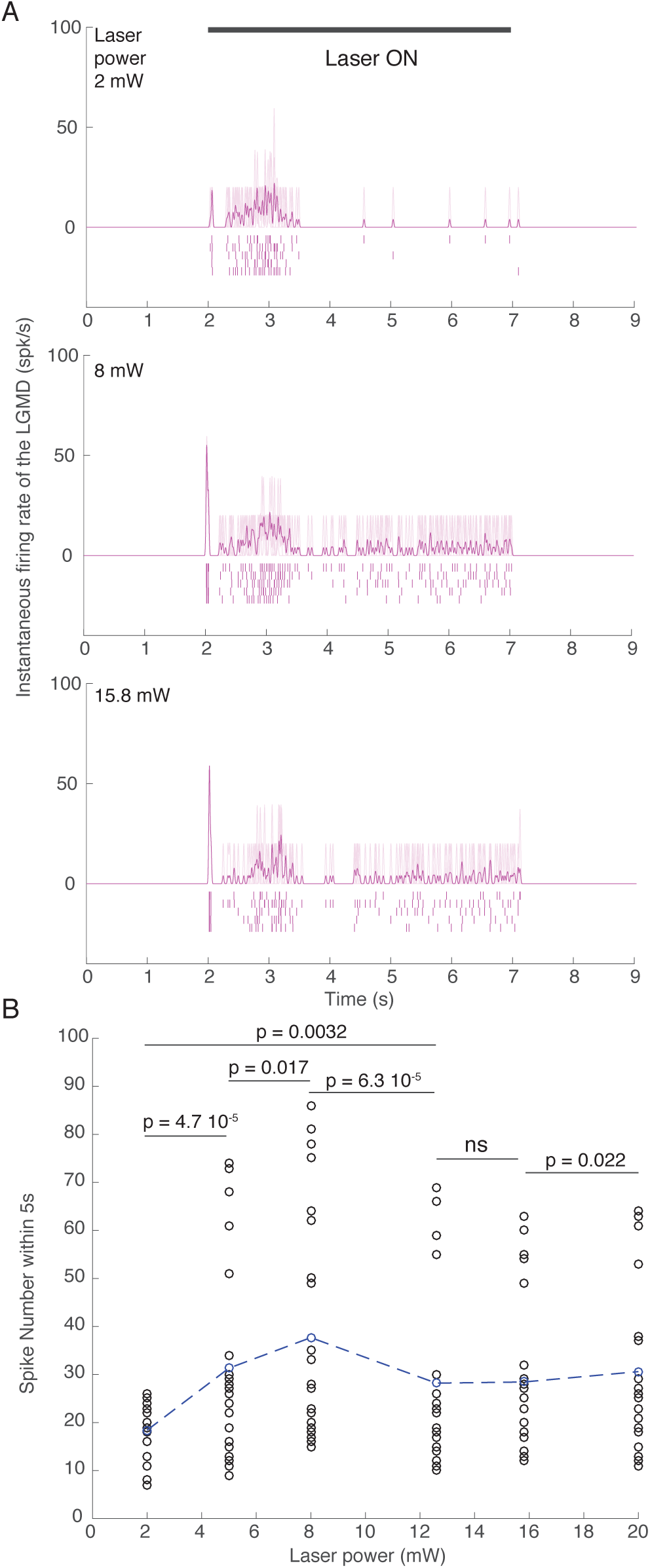
The firing of the LGMD increased and saturated in response to increasing laser power. **A**, Examples of the LGMD IFR in response to laser powers of 2, 8 and 16 mW (from top to bottom; one locust). Laser stimulation timing is shown above the top panel (duration: 5 s). Each panel shows the averaged LGMD IFR (dark purple) from 5 trials (light purple). The rasters below the LGMD IFR are the LGMD spikes from the 5 trials (p = 0.0312 and 0.125 between groups with laser power at 2 vs. 8 and 8 vs. 16 mW by a one-sided WSRT). **B**, The number of LGMD spike evoked by laser stimulation increased and saturated with increasing power. Blue circles indicate the mean value in each group. A Kruskal-Wallis test was used to evaluate the effect of laser power across groups (p=0.013). A post-hoc signed rank test was used to evaluate the difference between two groups (p values on panel; ns, no significant difference).

### Narrow laser beam produces less LGMD activation

When we varied the diameter of the optic fibers used to deliver the stimulus to the optic lobe from 10 to 200 μm we found that the illuminated region did not change much, always being ∼1 mm^2^. To further spatially restrict the number of neurons activated by the laser stimulus, we replaced the optic fiber with a specialized custom laser probe with an exit beam diameter of 10 μm (Fig. 5A; Methods). As demonstrated in Fig. 5B, illumination transmitted by this laser probe triggered EPSPs but no spiking in the LGMD, except for the spikes evoked by the on and off stimulation caused by the laser light onset and offset. The average LGMD’s V_m_ during laser probe illumination was significantly depolarized 0.5 ± 0.3 mV and 0.2 ± 0.3 mV in the first 2 s and over the whole duration of laser illumination. These results indicate that the modified laser probe is suitable for restricting the number of neurons activated by laser light.

## Discussion

In this study, we demonstrated that SFV A7(74) drove ChopWR-Venus expression on the cell membrane of medullary neurons in the locust optic lobe. Laser illumination increased the firing rate of the medullary neurons expressing ChopWR-Venus and triggered the firing of a downstream lobula neuron, the LGMD, which plays a key role in the locust visual collision-detection circuit. SFV A7(74) mediated highly efficient expression of ChopWR-Venus and led to the labelling of many neurons in the region surrounding the injection site. These findings provide a way to express genes of interest in locust neurons and to modulate neuronal activity using optogenetics. Besides optogenetics, GCaMP calcium indicators (Akerboom et al., 2012) are future candidates for expression in locust medullary neurons via SFV A7(74) transfection. These tools will help identify anatomically medullary neurons and study their roles in specific visual processing tasks through characterization of their calcium responses to different visual stimuli.

The responses immediately after the laser onset and offsets (arrowheads in Fig. 2; initial spikes in Figs. 3 and 4) were likely caused, in part, by photoreceptor activation from scattered laser light hitting the back of the eye. In experiments without viral transfection, no prolonged change in LGMD activity occurred in response to laser stimulation (Fig. 2E, bottom). The prolonged activation of medullary neurons and the LGMD during laser illumination were not attributable to the activation of photoreceptors, and therefore are believed to be solely due to the light-gated ChopWR current influx into the medullary neurons.

Because the large number of medullary neurons expressing ChopWR-Venus contained a mixture of both excitatory and inhibitory neurons, it was hard to precisely control neuronal activation of the LGMD through optogenetic stimulation. Reducing the area of laser illumination is one way to get more specific activation. We tested a specialized custom laser probe that minimizes the size of the laser beam and thus limits the number neurons activated (Fig. 5). In this configuration, only EPSPs but no spikes were evoked in the LGMD by optogenetic stimulation. However, a cell-type specific pattern of neural activation could not be achieved with currently available tools. In genetic model systems such as mice and fruit flies there are ways to generate cell-type specific gene expression. In mice, for example, the Cre-LoxP system drives cell-type specific expression through defined promoters (Sauer, 1998). In flies, the Gal4-UAS binary system mentioned above achieves the same goal (Busson and Pret, 2007). For transient gene expression in locusts, it would also be desirable to target genes to specific cell types. However, this is not yet feasible due to lack of identification of cell-type specific promoters and of transgenic locust lines expressing an effector gene under their control.

Yet, other possibilities to target gene expression in specific cell types exist. Micro RNAs (miRNAs) are small noncoding RNAs involved in posttranscriptional regulation of gene expression (Obernosterer et al., 2006). Recently, miRNAs have been applied to de-target gene expression when using a SFV-derived oncolytic virus to treat tumors such as glioblastoma (Ramachandran et al., 2017; Ylösmäki et al., 2013). The working principle of miRNAs is that by integrating the complementary sequence of a miRNA in the viral genome downstream of the viral subgenomic promoter, the miRNAs expressed in specific cells can identify the complementary sequence and cause the degradation of viral mRNA. This in turn reduces the expression of viral proteins in those cells. Wild-type SFV is naturally neurotropic (e.g., Ehrengruber et al., 1999). So, to protect neurons from SFV based cancer virotherapy, the neuron-specific miRNAs, miR124, miR125, and miR134 were inserted into the SFV4 vector genome. This resulted in attenuated neurovirulence in cultured neurons, astrocytes, and oligodendrocytes, and it also attenuated neurovirulence in adult mice, but the modified virus retained its replication ability in murine neural stem cells where the expression of these miRNAs is low (Ramachandran et al., 2017; Ylösmäki et al., 2013).

Interestingly, abundant miRNAs have been identified in tissues of a species closely related to that studied here, *Locusta migratoria*, including the pronotum, testes, antennae, fat bodies, and brains (Wang et al., 2015). Homology searches indicated that tissue-specific miRNAs were also lineage-specific and that many of them were specifically expressed in the brain. Thus, provided information becomes available in the future on the expression pattern of the miRNAs in specific cell populations, such as excitatory or inhibitory neurons, one could add their complementary sequence to SFV plasmids and obtain specific gene expression in the locust.

Injecting a viral vector containing a gene of interest in insects will only produce transient expression. Creating a transgenic line would be optimal to get stable foreign gene expression. Although there are no reported transgenic locusts, transgenic houseflies (Hediger et al., 2001), silkworms (Tamura et al., 2000), ants (Trible et al., 2017), honeybees (Schulte et al., 2014), and crickets (Nakamura et al., 2010) have been created using a transposon *piggyBac*-derived vector. In the locust, one recently identified miRNA precursor is likely a transposable element (TE) from a long-interspersed element family (Wang et al., 2015). One could thus try this putative locust specific transposable element or use the transposon *piggyBac*-derived vector described by Nakamura et al. (2010) to generate germ transformation.

## Acknowledgements

We thank Dr. Hiromu Yawo and Dr. Toru Ishizuka for providing pChopWR-Venus. We also thank Dr. Emma Victoria Jones and Dr. Keith Murai for providing the plasmids pENTR2B and pScaA7-RFA, and Dr. Alan L. Goldin for pSFV-Helper2.

**Authors’ contributions**
H.W., M.U.E., J.R., M.L.R. and F.G. designed the experiments; E.S. fabricated the laser probe; H.W. performed the experiments with the assistance of R.B.D and J.R.; H.W. carried out the data analysis with the assistance of R.B.D.; H.W. and F.G. co-wrote the paper.

